# Influenza Interaction with Cocirculating Pathogens, and Its Impact on Surveillance, Pathogenesis and Epidemic Profile: A Key Role for Mathematical Modeling

**DOI:** 10.1101/203265

**Authors:** Lulla Opatowski, Marc Baguelin, Rosalind M Eggo

**Author notes:** These authors contributed equally to the work (Lulla Opatowski, Rosalind M Eggo). Corresponding author (LO). Author Contributions:* LO, RME, MB conceived this review. LO and RME wrote the first draft. All authors contributed to the manuscript and approved the final version.

## Abstract

Evidence is mounting that influenza virus, a major contributor to the global disease burden, interacts with other pathogens infecting the human respiratory tract. Taking into account interactions with other pathogens may be critical to determining the real influenza burden and the full impact of public health policies targeting influenza. That necessity is particularly true for mathematical modeling studies, which have become critical in public health decision-making, despite their usually focusing on lone influenza virus acquisition and infection, thereby making broad oversimplifications regarding pathogen ecology. Herein, we review evidence of influenza virus interaction with bacteria and viruses, and the modeling studies that incorporated some of these. Despite the many studies examining possible associations between influenza and *Streptococcus pneumoniae, Staphylococcus aureus, Haemophilus influenzae, Neisseria meningitides*, respiratory syncytial virus, human rhinoviruses, human parainfluenza viruses, etc., very few mathematical models have integrated other pathogens alongside influenza. A notable exception is the recent modeling of the pneumococcus-influenza interaction, which highlighted potential influenza-related increased pneumococcal transmission and pathogenicity. That example demonstrates the power of dynamic modeling as an approach to test biological hypotheses concerning interaction mechanisms and estimate the strength of those interactions. We explore how different interference mechanisms may lead to unexpected incidence trends and misinterpretations. Using simple transmission models, we illustrate how existing interactions might impact public health surveillance systems and demonstrate that the development of multipathogen models is essential to assess the true public health burden of influenza, and help improve planning and evaluation of control measures. Finally, we identify the public health needs, surveillance, modeling and biological challenges, and propose avenues of research for the coming years.

**Author Summary:** Influenza is a major pathogen responsible for important morbidity and mortality burdens worldwide. Mathematical models of influenza virus acquisition have been critical to understanding its epidemiology and planning public health strategies of infection control. It is increasingly clear that microbes do not act in isolation but potentially interact within the host. Hence, studying influenza alone may lead to masking effects or misunderstanding information on its transmission and severity. Herein, we review the literature on bacterial and viral species that interact with the influenza virus, interaction mechanisms, and mathematical modeling studies integrating interactions. We report evidence that, beyond the classic secondary bacterial infections, many pathogenic bacteria and viruses probably interact with influenza. Public health relevance of pathogen interactions is detailed, showing how potential misreading or a narrow outlook might lead to mistaken public health decisionmaking. We describe the role of mechanistic transmission models in investigating this complex system and obtaining insight into interactions between influenza and other pathogens. Finally, we highlight benefits and challenges in modeling, and speculate on new opportunities made possible by taking a broader view: including basic science, clinical relevance and public health.

## Introduction

Influenza virus is a major contributor to the global disease burden, and exploration of its pathogenesis, epidemiology, and evolution has occupied generations of scientists. Its complex seasonality, antigenic drift of surface proteins, wide spectrum of severity, and capacity to cross species and cause epidemics or pandemics are all characteristics that make the virus so difficult to control [1].

The human respiratory tract is an important reservoir of bacteria, fungi, viruses, bacteriophages, archaea and eukaryotes [2], harboring diverse communities of commensal, opportunistic and pathogenic microorganisms. It has been suggested that some of these species enter into non-neutral relationships [3], including competition for resources, synergism with the host immune system, or physiological modifications that alter the normal colonization or infection processes. The contribution of these phenomena to the influenza-infection burden is largely unknown.

In terms of public health, what is understood concerning influenza transmission or severity may therefore be incomplete or misguided due to ignorance of the effect of interacting pathogens. On one hand, large-scale influenza vaccination programs may unexpectedly impact other infections due to an indirect rise or fall in the risk of contracting them in a pool of influenza-infected individuals [4]. For example, if influenza outcompetes another virus and holds it at bay, an influenza vaccination program could result in an upsurge in the competitor. On the other hand, the introduction of measures to control bacterial infections (e.g. pneumococcal vaccines) may indirectly and positively impact the influenza disease burden as secondary bacterial pneumonia are associated with severe outcomes of influenza.

Seasonal influenza generates a huge burden each year during the wintertime in temperature regions and with more complex seasonal patterns in tropical regions [5]. However, influenza pandemics frequently occur outside of the usual season, and generate an unpredictable and often large burden in morbidity, mortality, and cost [6,7], mostly due to devastating role of secondary bacterial infections [8,9]. This out-of-season circulation of pandemic strains takes place in different climatic and ecological milieus than seasonal strains, and therefore pandemic strains may interact with different coinfectors. It is therefore critically important to pandemic preparedness to understand competitive and synergistic relationships. Considering influenza in a context of interactions with other environmental species, both at the individual level from a clinical perspective, or at a population level from an epidemiological perspective, is vital to improve our understanding and control of virus transmission and the risk of developing disease on infection.

Mathematical modeling has been a key tool in infectious diseases for many years because it links the transmission of an infection from person-to-person to the dynamics of the infection at a population level [10]. Models allow researchers to probe the complex intricacies of transmission, and play forward the effects on an individual to see the impact on population level infection dynamics. Researchers can therefore easily create counterfactuals: “what if” scenarios; where vaccination rates, contact patterns, health behaviors, or any number of other factors, are different.

Models of influenza virus transmission have proved very useful in expanding knowledge of influenza biology, evolution and epidemiology. For example, models of evolutionary change and immunity aim to predict the dominant strain of influenza in the coming season [11]. Spatially explicit models have convincingly linked commuting movements to the spread of influenza in the USA [12]. Models have also been crucial to public health, contributing to the optimization of control strategies, including use of vaccines and antivirals [13–19]. As the modeling field has developed, there has been effort to improve realism by incorporating heterogeneity in human contact patterns, age-related susceptibility, cross immunity after previous infections [18,20–23], and the potential effect of environmental variables on transmission [12,24]. Notably the vast majority of modeling work has neglected the microbial environment. Most mathematical and computational models of influenza are focused on single or sequential influenza-only infections and have broadly simplified pathogen ecology. For example, models have not been exploited to estimate the indirect effect of seasonal influenza vaccination on the incidence of severe bacterial infections in the elderly. Further, despite secondary bacterial infections being recognized as an important cause of mortality, modeling used to plan vaccine interventions during the pandemic in the considered influenza transmission alone [25].

The authors of relatively recent literature reviews gathered biological and epidemiological evidence for interactions between influenza virus and respiratory bacteria or viruses [3,26,27], but did not consider mechanistic transmission models. Mathematical models can be used to investigate mechanisms of interaction, and visualize the pathological and epidemiological patterns that result from them. Model outputs can be compared with or fitted to data, thereby enabling estimation of both the probability of such interactions, and the strength of the interaction. Estimation can be made across geographic regions (eg. winter seasonal vs year-round-transmission), for different virus subtypes (e.g. seasonal vs pandemic), and in different age groups (e.g. infants vs elderly). Computational and mathematical models to study influenza with other respiratory pathogens are currently underutilized.

In this review, we report evidence of influenza interaction with other pathogens and systematically review the modeling studies on influenza coinfection. We address how different interference mechanisms might lead to unexpected epidemiological patterns and misinterpretations. Finally, we identify public health needs, modeling and biological challenges, and propose avenues of research for the coming years.

## Mechanisms of interaction

Here, “interaction” refers to any process by which infection caused by one pathogen affects the probability, timing, or natural history of infection by another. This process includes a wide range of mechanisms that can involve direct connections between the two pathogens, e.g. at the cellular level, or indirect interactions through an intermediate factor that influences the other. The indirect consequences of these interactions are described later. For influenza virus, interactions with bacterial or viral species can occur at several scales (Fig 1). Interacting pathogens may have two distinct profiles: natural human commensals, usually bacteria, which cause mainly asymptomatic carriage or mild symptoms often for long durations of weeks to months; or epidemic pathogens causing infection for shorter durations, from a few days to a few weeks. These two distinct epidemic profiles potentially involve different modes of interaction and lead to different levels of consequences. Here we detail proven and potential interaction mechanisms (Fig 1).

**Fig 1.**
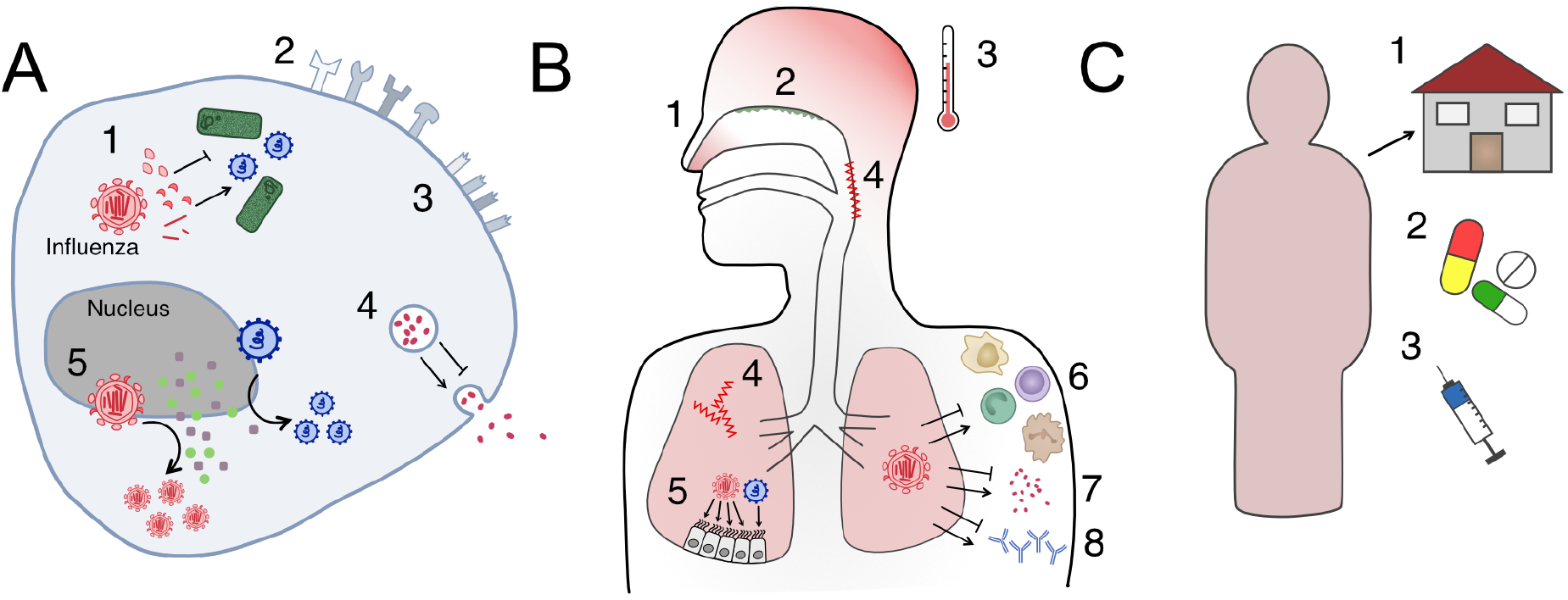
Influenza interactions with other pathogens occur within-host or at the population level. Each interaction could either inhibit or enhance coinfection, depending on the combination of pathogens. **A**) Cellular level interactions: 1. direct interactions between viral products; 2. altered receptor presentation; 3. cell damage, e.g. its surface receptors; 4. modification of release of immune-system mediators; 5. competition for host resources among influenza and other pathogens. **B**) Host-level interactions: 1. change of transmissibility due to symptoms; 2. individual variation in commensal microbiota; 3. effect of symptomatic responses to infection; 4. tissue damage, e.g. in the nasopharynx or lung; 5. competition for host resources, e.g. target cells for infection; 6. immune-cell-mediated interaction; 7. immune signaling-mediated interaction; 8. antibody-mediated interaction. **C**) Population-level interaction: 1. behavioral responses to disease; 2. medication use; 3. vaccination behavior.

### Within-Host Interactions

At the cellular level, interactions involve both direct and indirect mechanisms. First, influenza genes or gene products can enhance or inhibit the replication of other viruses or potential infection by bacteria [28] by direct interaction with pathogen proteins or nucleic acids. Further, indirect competition for host resources can occur, when pathogens compete for target cells, receptors, or cellular products required for replication, thus preventing or inhibiting superinfection by a secondary virus. Influenza-infected cells may also release cell-signaling molecules that could increase or decrease the probability of coinfection.

During infection influenza virus impairs innate and adaptive host defenses [29,30]. Mechanisms include altered neutrophil recruitment and function, leading to defective bacterial clearance, diminished production of alveolar macrophages [31] and inhibition of T-cell-mediated immunity [30]. Infection with a second virus could be modulated similarly, e.g. by the production of cross-reactive antibodies or cell-mediated immunity that prevents or facilitates this infection. Physiological changes induced by the host response to infection may have ecological consequences. For instance, lung-tissue damage [31] and the induction of type-1 interferon signaling were shown to promote bacterial colonization [30], and broadly inhibit virus replication [32]. Damage to lung cells caused by influenza infection, such as influenza neuraminidase stripping sialic acids from the cell surface, amplifies bacterial adherence and invasion [26], and could potentially change the likelihood of infection by another virus. Symptomatic responses to infection, like fever, have also been shown to act as “danger signals” for bacteria, e.g. meningococci, which react by enhancing bacterial defenses against human immune cells [33]. In contrast, fever may diminish viral replication rate, thereby lowering the probability of coinfection. Not all interactions depend on what happens after the influenza infection: the pre-infection respiratory flora of infected individuals may partially account for the variability of influenza infection severity and outcome [27]. For example, in animal studies, *S. aureus* colonization was shown to trigger viral load rebounds and reduced virus clearance [34–36].

### Population-Level Interactions

Behavioral responses to influenza infection can also indirectly impact the transmission of bacteria or other viruses. On one hand, people with severe influenza symptoms are likely to stay home, modifying their contact patterns, and making acquisition of second infections unlikely [37,38]. On the other hand, individuals with milder symptoms might maintain their regular activities, which could increase bacterial transmission to other individuals (as observed for tuberculosis [39]) or increase the chance of acquiring a second infection. Person-to-person variation in care seeking and medication use, such as of antivirals, antibiotics, antipyretics or vaccine(s) can also influence the risk of coinfection. For example, use of the pneumococcal conjugate vaccine has decreased carriage of the bacteria in some contexts [40,41], which may lead to a decreased chance of observing influenza-pneumococcal coinfections. Public health policies, such as vaccination or pharmaceutical recommendations targeting one pathogen may also indirectly impact another (Fig 2). Further discussion of the effect of public health interventions is given in the population impact section.

**Fig 2.**
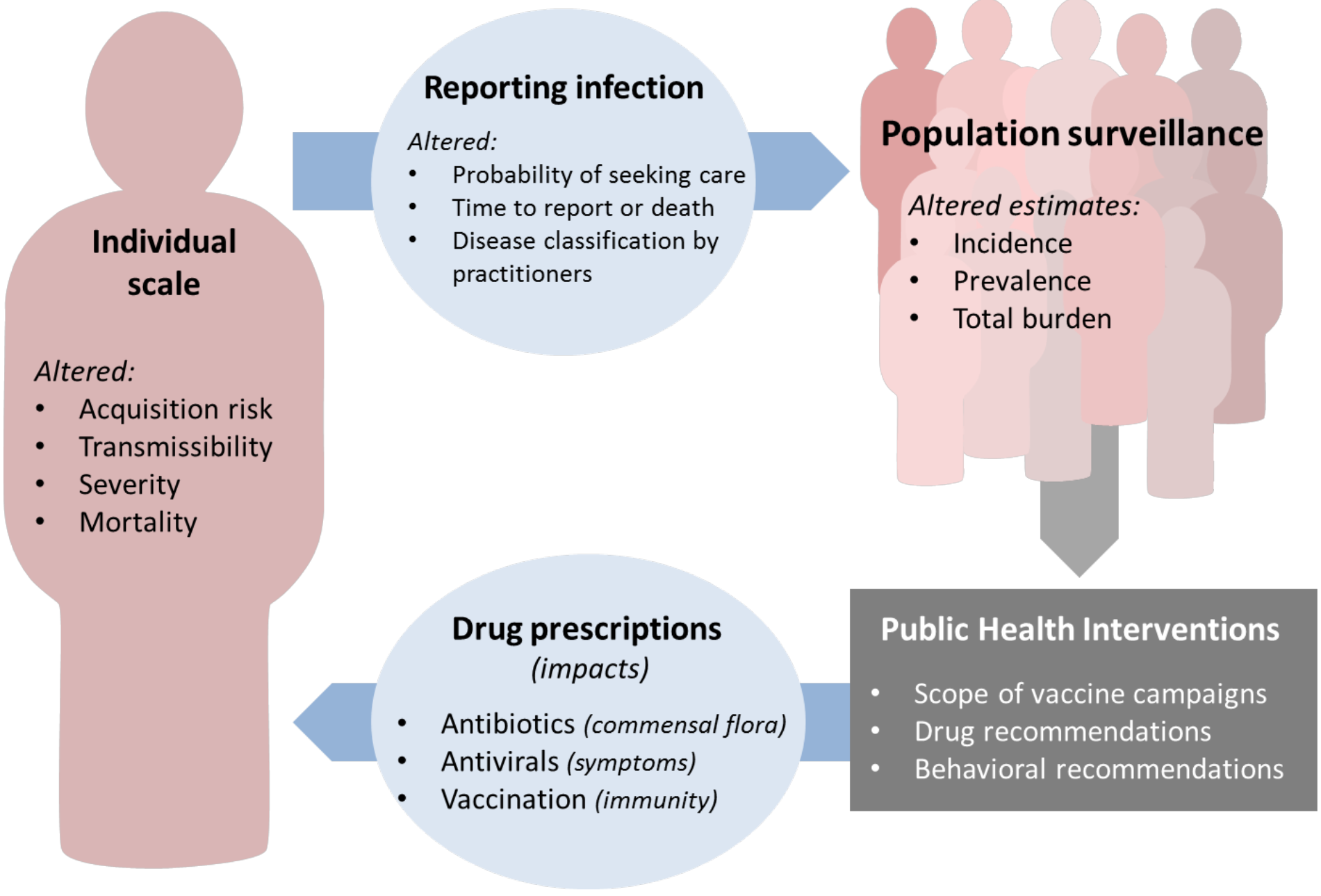
Cycle of factors affected by non-neutral interactions at the individual level and their impact on influenza surveillance, treatment, prevention and control. Factors that affect coinfection on an individual scale can feed forward to an effect on population surveillance through their effects on the reporting of infection. Decisions on public health interventions are made in response to population-level data. These interventions then take effect at the individual level, to give a feedback loop both generated and impacted by effects of coinfection.

## Evidence of interaction

Several literature reviews described evidence of interactions between influenza virus and other respiratory bacterial or viral pathogens [3,26]. Here we describe the viruses and bacteria that potentially interact with influenza by gathering recent findings from laboratory and epidemiological studies for bacteria and viruses (Supplementary Section A).

### Influenza-bacterial interactions

Experimental results suggest that most of the pathogenic and commensal bacteria present in the nasopharynx may directly or indirectly interfere with influenza infection during host colonization or infection (Table 1). The best-studied influenza-bacterial interaction is with *Streptococcus pneumoniae* [3]. Influenza is thought to increase bacterial adherence and facilitate the progression from carriage to severe disease [27,42], although evidence from population studies is not so clear-cut [43–46]. Influenza was also shown to impair methicillin-resistant *Staphylococcus aureus* (MRSA) clearance in coinfected mice, thereby increasing their susceptibility to MRSA infection [47]. Similarly in mice, increased severity of *Haemophilus influenzae* induced by influenza virus was suggested based on experiments of sequential infection with sublethal influenza then *H. influenzae* doses [48]. Notably, ecological studies revealed a positive association between influenza and *Neisseria meningitides* incidence [49] and those of *in vitro* studies suggested that direct interaction between influenza A neuraminidase and the *N. meningitidis* capsule enhanced bacterial adhesion to cultured epithelial cells [50]. Lastly, in patients with pulmonary tuberculosis there is evidence of increased risk of severe outcomes on influenza infection [51]. This finding was supported by experiments in mice [52] which demonstrated that *Mycobacterium tuberculosis* and influenza coinfected mice mounted weaker immune responses specific to *Mycobacterium bovis* Bacillus Calmette-Guerin (BCG) in the lungs compared to mice infected with BCG alone.

**Table 1.** Bacteria whose colonization or Infection course may be affected by interaction with influenza.

| Bacterial species | Study system | Effect | Illustrative publications |
| --- | --- | --- | --- |
| *S. pneumoniae* | Animal | Synergistic/Facilitating | Smith 2013 [90]; Wolf 2014 [130]; Siegel 2014 [131]; McCullers 2010 [132]; Ghoneim 2013 [31]; Peltola 2006 [133]; Walters 2016 [134]; Nakamura 2011 [135]; |
|  | Human | Synergistic/Facilitating | Walter 2010 [136]; Nelson 2012 [137]; Opatowski 2013 [95]; Shrestha 2013 [100]; Weinberger 2013 [86]; Jansen 2008 [138]; Kuster 2011 [139]; Nicoli 2013 [85]; Ampofo 2008 [140]; Grabowska 2006 [141]; Murdoch 2008 [142]; Edwards 2011 [143]; Weinberger 2014 [144]; Grijalva 2014 [145]; |
|  |  | Neutral / Unclear | Kim et al 1996 [44]; Watson 2006 [45]; Toschke 2008 [46]; Zhou 2012 [146]; Damasio 2015 [147]; Hendricks 2017 [148] |
| *S. aureus* | In vitro and animal | Synergistic/Facilitating | Niemann 2012 [149]; Davison 1982 [150]; Tashiro 1987 [36]; Zhang 1996 [151]; Chertow 2016 [152]; Sun 2014 [47]; Braun 2007 [34]; Iverson 2011 [153]; Robinson 2013 [154] |
|  | Human | Synergistic/Facilitating | Sherertz 1996 [155]; Hageman 2006 [156]; Finelli 2008 [157]; Reed 2009 [158] |
|  |  | Neutral | Kobayashi 2013 [159] |
| *H. influenzae* | Animal | Synergistic/Facilitating | Lee 2010 [48]; Michaels 1977 [160]; Bakaletz 1988 [161]; Francis 1945 [162] |
|  | Human | Synergistic/Facilitating | Morens 2008 [163] |
| *N. meningitidis* | In vitro and Animal | Synergistic/Facilitating | Rameix-Welti 2009 [50]; Loh 2013 [33] |
|  |  | Neutral | Read 1999 [164] |
|  | Human | Synergistic/Facilitating | Cartwright 1991 [165]; Hubert 1992 [49]; Jacobs 2014 [166]; Brundage 2006 [167]; Jansen 2008 [138]; Jacobs 2014 [166]; Makras 2001 [168] |
| *M. tuberculosis* | Animal | Synergistic/Facilitating | Florido 2015 [169]; Florido 2013[52]; Volkert 1947 [170]; Redford 2014 [171] |
|  | Human | Synergistic/Facilitating | Walaza 2015 [172]; Oei 2012[173]; Noymer 2011 [174]; Noymer 2009 [175]; Zurcher 2016 [176] |
|  |  | Neutral | Roth 2013[177] |
| *S. pyogenes* | Animal | Synergistic/Facilitating | Klonoski 2014 [178]; Okamoto 2003 [179]; Okamoto 2004 [180]; Hafez 2010 [181] |
|  | Human | Synergistic/Facilitating | Scaber 2011 [182]; Zakikhany 2011 [104]; Tasher 2011 [183] |
|  |  | Neutral | Tamayo 2016 [184] |

### Virus-virus interactions

Within its family, influenza interacts between types (A and B), subtypes (e.g. H3N2, H1N1) and strains. Competitive exclusion due to homologous immunity is widely accepted [53,54], and has been applied extensively in models of influenza-strain coexistence [55,56]. Antigenic change (measured through antigenic distance) occurs constantly in influenza, strongly indicating that the virus escapes from immunity resulting from prior infection by genetic change [57]. There is mounting evidence that the first influenza infection is important, and may affect severity of future infections [58–60]. Some evidence also supports the finding that influenza can interact with other influenza viruses, and non-influenza respiratory viruses via non-specific immunity following infection [61,62].

For non-influenza interactions, many viruses are suspected of interfering with influenza virus acquisition, based on different types of studies (Table 2). During the 2009 influenza pandemic, Casalegno et al. reported that, in France, the second pandemic wave was delayed due to the September rhinovirus epidemic [63], even though the shift was not observed in other countries [64,65]. Coinfection by the two viruses might also enhance disease severity for individuals [66–68], although evidence is discordant [69–71]. Similarly, competitive interaction with RSV has been posited for many years [72,73], and some evidence was found for delayed RSV epidemics due to the second wave of the 2009 pandemic in France [74] and tropical regions [75,76]. There is discrepancy in the findings of interaction between influenza and RSV, with most studies finding increased severity [71,77,78] whereas others found no effect [66] and some found less severity [79]. Competitive interaction with parainfluenza viruses was also inferred, based on less frequent coinfection pairs than expected [80], but that observation is not consistent across studies [81–83]. In terms of severity, parainfluenza and influenza coinfection is usually more severe than influenza alone [66,68,84] but not always [69,70].

**Table 2.** Viruses that may be affected by interaction with influenza.

| **Virus** | **Study system** | **Effect** | **Illustrative publications** |
| --- | --- | --- | --- |
| RSV | Population incidence | Competitive | Anestad 2007 [185]; Anestad 2009 [186]; Casalegno 2010 [74]; , Anestad 1987 [187]; Yang 2012 [76]; Nishimura 2005 [188]; Glezen 1980 [73]; Pascalis 2012 [80]; Yang 2015 [64]; van Asten 2016 [189]; Meningher 2014 [190]; Velasco-Hernandez 2015 [110] |
|  |  | Neutral | Navarro-Mari 2012 [65] |
|  | Coinfection detection | Competitive | Greer 2009 [81]; Martin 2013 [191] |
|  | Laboratory investigation | Competitive | Shinjoh 2000 [192] |
| Rhinovirus | Population incidence | Competitive | Casalegno 2010 [63]; Casalegno 2010 [74], Pascalis 2012 [80]; Linde 2009 [193]; Anestad and Nordbo [194]; Cowling 2012 [62]; Yang 2015 [64] |
|  |  | Neutral | Yang 2012 [76] ; Navarro-Mari 2012 [65]; van Asten 2016 [189] |
|  | Coinfection detection | Competitive | Tanner 2012 [195]; Mackay 2013 [196]; Nisi 2010 [83]; Greer 2009 [81]; Martin 2013 [191] |
|  | Laboratory investigation | Competitive | Pinky and Dobrovolny 2016 [105] |
| Influenza | Population incidence | Competitive | van Asten 2016 [189] |
|  | Coinfection detection | Competitive | Nisii 2010 [83]; Sonoguchi 1985 [53] |
|  | Laboratory studies | Competitive | Easton 2011 [197]; Laurie 2015 [54] |
| HPIV | Population incidence | Competitive | Yang 2012 [64]; Anestad 1987 [187] ;Yang 2015 [64] |
|  |  | Neutral | Mak 2012 [75] |
|  | Coinfection detection | Competitive | Pascalis 2012 [80] |
|  |  | Neutral | Murphy 1975 [82]; Nisii 2010 [83]; Greer 2009 [81]; Martin 2013 [191] |
|  | Laboratory investigation | Synergistic/<br>Facilitating | Goto 2016 [198] |

The general pattern is that bacteria tend to synergize with influenza, often boosting transmission of either pathogen, or increasing invasion of the bacteria following influenza infection. It is not always clear if this is a true synergy - where both pathogens benefit - or rather that influenza facilitates bacterial invasion. In contrast, viral pathogens tend to form competitive interactions with influenza, although whether these are direct, specific interactions with particular other viruses, or the result of an “early advantage” to the first infector, remains unclear. This pattern may occur because of the differing natural histories of bacteria and viruses; where the former tends to infect hosts for long time periods, and the latter has shorter infections, more similar to the natural history of influenza itself.

## Impact of Interactions at the Population Level

Although coinfections occur at the host level, their consequences are far-reaching (Fig 2). Coinfection may alter the natural history, severity or timing of illness in an individual, and thereby modify the morbidity, healthcare-seeking behavior and treatment of that individual. Heterogeneity in these can affect the probability of, and timing of, reporting disease, thereby transferring the effect from individual hosts to the population level.

Development and implementation of public health policies rely on analyses of population surveillance data on influenza epidemics and burden. Policies then generate changes in medical interventions at the population level, e.g. change in vaccination rates, or at the individual level, e.g. recommendations for antibiotics or antivirals in certain groups. These public health interventions then have their own impacts on the dynamics of pathogens and coinfections. Therefore as coinfections may alter surveillance data, and policies based on evidence from surveillance data may alter coinfection or interference risk, there is a complex cycle of dependence, which highlights the difficulty—as well as the potential importance—of assessing the impact of coinfections (Fig 2).

To date, most of the published quantitative analyses of interactions rely on statistical association between incident cases of influenza-like-illness (ILI) and other infections based on regression and correlation analyses [85,86]. A major methodological challenge of detecting interactions is that significant correlation between epidemics of two pathogens in surveillance data may result from either a true biological direct or indirect interaction, or may be confounding as a result of the two pathogens sharing common ecological conditions (e.g. cold weather). Regression models do not formalize the transmission process or biological mechanism of interaction. Instead, they describe simple functional links between, for example, the incidence time series, onset or peak time, or epidemic magnitude or severity. They provide correlations between reported time series at different time lags and are useful tools to detect strong signals of associations, however, despite their apparently simple formulation, regression models “assume” strong statistical hypotheses based on the shape of the data and the association [87]. These coarse methods cannot disentangle intricate interaction mechanisms and, furthermore, the lack of mechanistic formulation of transmission and interaction hinders quantification of interaction strength, and prevents easily interpretable predictions that can be used for public health decision-making.

Due to the complex phenomena involved and many feedback interactions, mechanistic models are needed to dissect the cause-and-effect of the different components [88] (Box 1). The role of modeling is two-fold: first, mathematical modeling provides a common language to integrate heterogeneous mechanisms and test competitive hypotheses. By doing so, models contribute to building basic knowledge about infection processes. Second, modeling enables assessment of potential intervention scenarios by predicting their impact.

For these reasons, public health interventions based on modeling of infectious diseases have become informative and effective. For example, in the UK, a transmission model fitted to a vast range of ILI and influenza-surveillance data demonstrated that vaccinating children against influenza will have the same protective effect on people over 65 years old, as vaccinating those individuals [89]. This outcome is a consequence of the diminished community transmission that results from reducing infections in children. Such an impact would be impossible to identify without mechanistic models. Box 2 summarizes the benefits of mathematical models.

## Models of influenza interactions

Despite mounting evidence of influenza-bacteria interactions and the concurrent increasing use of dynamic modeling to study infectious diseases in recent decades, influenza interactions have rarely been modeled. Interestingly, previous literature reviews describing evidence of interactions between influenza virus and other respiratory bacterial or viral pathogens [3,26] neglected mathematical models which, despite their limited number, provide insight into mechanisms of interaction and their consequences. We have systematically reviewed the literature for models incorporating influenza with bacteria or non-influenza viruses (Supplementary Section A).

### Influenza-bacterial interaction

The only influenza-bacterium interaction that has been integrated into mathematical modeling studies is the influenza-pneumococcus system, both within-host and at the population level.

Several dynamic models of coinfection at the cellular level were proposed relatively recently [90–92,93]. In a study combining modeling and empirical data from mice coinfected with two different influenza viruses and two pneumococcus strains, Smith et al. assessed the likelihood of different immunological interaction mechanisms [90]. They found a role of macrophage dysfunction leading to an increase of bacterial titers and increased virus release during coinfections [90]. However, their results suggest that coinfection-induced increase of bacterial adherence and of infected cell death were not very likely. Shrestha et al. used an immune-mediated model of the virus-bacterium interaction in the lungs to specifically quantify interaction timing and intensity [91]. They assumed that the efficiency of alveolar macrophages, which are a critical component of host immunity against bacterial infections, was reduced by viral infection and tested the impact of inoculum size, time of bacterial invasion after influenza infection, and the potential impact of antiviral administration. The model predicted that enhanced susceptibility to invasion would be observed 4-6 days after influenza infection, suggesting that early antiviral administration after influenza infection (<4 days) could prevent invasive pneumococcal disease. Smith & Smith modeled a nonlinear initial dose threshold, below which bacteria (pneumococcus) declined and above which bacteria increased. They showed that this threshold was dependent on the degree of virus-induced depletion of alveolar macrophages, using data from mice experiments. Because macrophage depletion varies through the course of influenza infection, this important finding may explain why risk of bacterial invasion also changes over the course of infection, with particularly low dose requirement in the first few days of infection [94]. In a follow up study, the same authors analyzed published data from influenza-pneumococcus co-infected mice treated with antiviral, antibiotic, or immune modulatory agents. They found that antivirals are more efficient at preventing secondary infection when used in the first two days of influenza infection; and also found an important benefit of immunotherapy, especially for low bacterial loads [92]. Lastly, in a within-host model, Boianelli and colleagues investigated the efficacy of different oseltamivir treatment regimens in influenza-pneumococcus coinfected individuals using parameters drawn from human and mouse studies. They found that increasing the dose of oseltamivir, but not duration of treatment, might increase both the antiviral and antibacterial efficacy [93].

There have been several population transmission models to assess influenza interactions with bacteria, and test hypotheses regarding the main mechanisms [95–99]. Analysis of pneumococcal meningitis and viral respiratory infections in France highlighted two important processes in colonized individuals: a virus-related increase in pneumococcal pathogenicity and enhanced transmissibility of bacteria to others [95]. Analysis of bacterial pneumonia from the USA concluded that influenza increased individual risk of pneumonia [96,100]. Recently, in a simulation study, Arduin *et al.* used a flexible individual-based model of influenza-bacterial interaction to assess the population consequences and associated burden of a range of pneumococcus-influenza interaction mechanisms [101]. Population dynamic models have also been used to test the public health impact of control measures [97–99]. Different strategies of antibiotic use (as treatment or prophylaxis) and of vaccination were assessed by modeling the dual transmission of pneumococcus and influenza [97]. When modeling a 1918-like pandemic, this model suggested that widespread antibiotic treatment of individuals with pneumonia would significantly lower mortality, whereas antibiotics in prophylaxis would effectively prevent pneumonia cases. A different model evaluated the benefit of vaccinating the UK population against pneumococcus in the context of pandemic influenza using different pandemic scenarios: 1918-like, 1957/68-like or 2009-like [98]. Those results indicated that pneumococcal vaccination would have a major impact only for pandemic with high case-fatality and secondary pneumococcal infections rates (e.g. the 1918-like), with less influence in other scenarios.

### Viral interaction

Influenza-influenza interactions predominate in models of two viruses, with limited investigation of influenza-RSV interactions, and no models of other viruses.

Within-host, several models of multi-strain influenza infections were proposed [102–104], especially examining the interval before the secondary infection. One model of RSV-influenza interaction at the cellular level explored the hypothesis of the viruses interacting through competition for resources within the cell [105]. This indirect competition was sufficient to explain the observed rate of virus replication. The model also explored how the speed of virus replication confers an advantage to the first infecting pathogen, and determined the “head start” on infection that the slower-replicating virus would require to maintain dominance.

Population models have been used extensively to examine the dynamics of influenza and multi-strain influenza systems (for a review see [106]) although many fewer studies examined multispecies systems. Because the influenza virus comprises two species, multiple subtypes, and potentially numerous strains of each, at any given time many viruses may be circulating, providing varying degrees of cross-protection after recovery, and sometimes with complex dynamics of within-species strain replacement due to genetic drift or reassortment. There is evidence of competition between strains, with some models requiring short periods of heterologous immunity after infection to create the ladder-like phylodynamic structure of influenza viruses [107], although recent studies could capture this feature without this mechanism [55]. One comprehensive early model tested four mechanisms of interaction between influenza types using data from Tecumseh, Michigan, but the data were insufficient to distinguish the mechanisms [108]. Influenza-influenza models must also account for the complex immune history of hosts, where there is mounting evidence that the timing of an individual’s influenza encounters, and especially the first infection, shapes their future response [58–60]. The methods for modelling influenza-influenza interactions should be extended into interactions with other viruses.

One model for pandemic influenza, in which coinfection with other respiratory pathogens leads to enhanced influenza transmission, was proposed to explain the multiple waves of the 1918 influenza pandemic in the UK [109]. A recent example of influenza and RSV crossspecies analysis, in a climatically driven model, provided some evidence that RSV dominates influenza, but the model was not explicitly fitted to data [110].

### Illustration from a simple model

To demonstrate how both synergistic and competitive interactions can be modeled, we used a simple transmission model and simulated the effect of interactions (Box 3, Figure 3, and Supplementary section B). We show how these interactions at the individual level can impact the epidemics at the population model. The “bacterial type” interaction firstly shows an increase in bacterial prevalence when influenza infection increases bacterial transmission, in a facilitative interaction. In a synergistic interaction, where coinfection increases transmission of both influenza and bacteria, prevalence of bacteria increases, and the epidemic of influenza has a quicker and higher peak. In the competitive interaction, progressively decreasing the probability that a second pathogen can infect an already-infected host causes the epidemic peaks to separate in time. It also decreases the peak size of the outcompeted pathogen, without altering the number of people infected in total (Figure 3).

**Fig 3.**
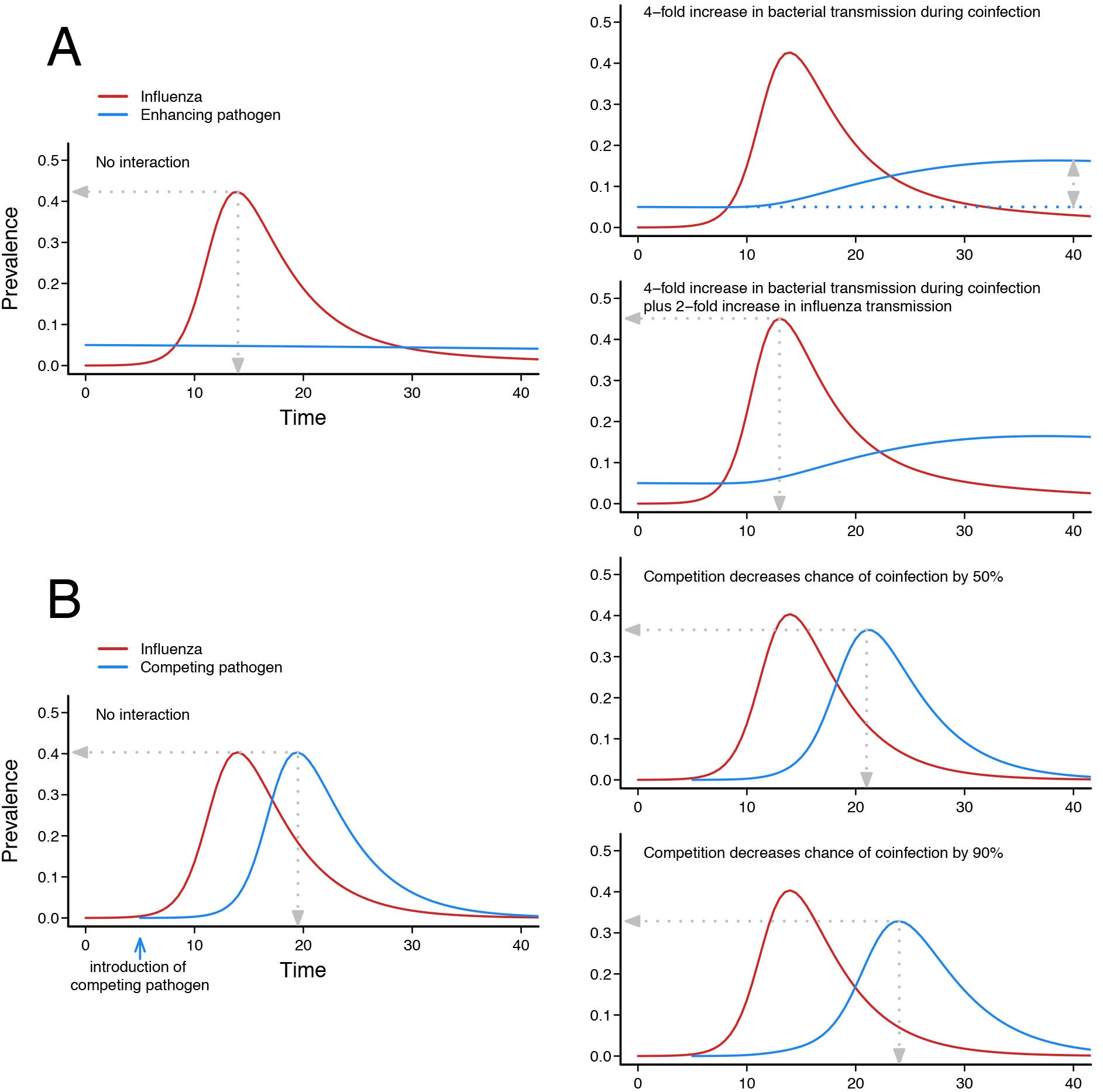
Example model outputs showing effect of synergistic and competitive interaction. Box 3 gives details on the model that produces these epidemic trajectories. A) In the baseline enhancing scenario, an endemic bacterial pathogen (blue) occurs at 5% prevalence. An influenza epidemic occurs with no interaction, and the bacterial prevalence does not change. If the presence of influenza coinfection increases bacterial transmissibility by 4 fold (*σ*_*1*_ = 4) then there is a transient rise in bacterial prevalence. If there is also an increase in influenza transmissibility during coinfection (*σ*_*1*_ = 4 and *σ*_*2*_ = 2) then there is also a higher and earlier influenza peak, as a result of coinfection. B) In the baseline competition scenario, the second epidemic pathogen is introduced later than influenza. The two pathogens have the same transmission characteristics (same *γ*, same β). If there is only a 50% chance of infection with pathogen 2 when individuals are infected with pathogen 1 (*δ*_*1*_ = 0.5) then the epidemic trajectory of pathogen 2 is lower and later. If competition is even stronger (*δ*_*1*_ = 0.1) so there is a 90% reduction in chance of coinfection, the profile of pathogen 2 is even further separated from pathogen 1. Computer code generating these trajectories is given in Supplementary File S1.

## Limits of the Current View

Historically, scientific and medical studies have tended to focus on host-pathogen interactions in an independent manner, by studying each pathogen alone. It is clear that the human host simultaneously encounters many microbes. Many respiratory viruses and bacteria have been linked to influenza epidemiology, based on *in vivo* evidence from individual and epidemiological studies. These non-neutral interactions, mostly facilitative for bacteria and competitive for viruses, probably have individual- and population-level effects on influenza pathogenicity, burden and potentially its epidemic profile. Furthermore, this is a complex system in which each host-pathogen or pathogen-pathogen interaction phenomenon may impact the others. Surprisingly however, such interactions remain poorly studied and, in particular, very few modeling studies have addressed these questions.

Mathematical models are crucial to guide public health decision-makers, who, for ethical or cost reasons, cannot conduct large-scale trials. Two examples of interventions based on modeling results and mobilizing important public resources are pandemic preparedness (stockpiling of antivirals, use of vaccine doses) [111] and national immunization programs [19]. Neglecting the cocirculating pathogens— i.e., adopting influenza tunnel vision—and the indirect impact of coinfections, may potentially affect the estimation of the risk associated with influenza infection and, consequently, the accuracy of model predictions. Interaction strength may also change from year-to-year, and depend on the circulating influenza strain. For evaluation of interventions, this neglect can lead to overestimation of the impact - if burden was measured without considering the changing landscape of coinfection in the population; or underestimation - if the effect of an intervention does not account for the potentially decreased burden of an interacting pathogen as a result of diminished influenza transmission. For all these reasons, we think that adopting a more holistic approach to modeling of respiratory pathogens will improve their surveillance and the strategy to control them.

## Opportunities

Considering influenza virus in its ecological context and its interactions as a cause of its associated morbidity and mortality should offer opportunities for prevention and treatment. On one hand, models can assess available vaccines in optimal combinations and for better-targeted populations. In addition to influenza vaccines that (partially) protect against virus acquisition, antibacterial vaccines are also critical. Pneumococcal vaccines have good efficacy against influenza-associated non-bacteremic pneumonias [112,113]. The 23-valent pneumococcal polysaccharide vaccine significantly lowered the risk of invasive pneumococcal disease and attributed mortality in the elderly [114]. Better understanding of possible influenza-pneumococcus interactions and integrating those into transmission models could potentially identify synergies between these vaccination programs, and optimize the use of both vaccines.

In addition, optimization of antibiotic and antiviral prescriptions should be considered. First, antibiotics have historically been used extensively to prevent secondary infections [115,116]. However, increasing rates of antibiotic resistance worldwide led to policies to mitigate antibiotic consumption, focusing particular attention on reducing antibiotic prescriptions for viral infections. Second, neuraminidase inhibitors were found to prevent some secondary bacterial pneumonias in animal models, human investigations and modeling studies, beyond the window in which they directly impact the influenza viral load [91,117,118]. Although antivirals may only modestly attenuate influenza symptoms, a body of evidence suggests they could avoid severe and economically important outcomes of influenza infection [118–121].

Accurate burden quantification is crucial to designing and implementing public health interventions against influenza. Focusing efforts to better understand these interactions is therefore critical, especially in the context of pandemic influenza, but also to plan for seasonal epidemics, by forecasting the onset and peak times, and estimating the expected burden. Deeper understanding of the ecology of the vast number of microorganisms that can contribute is needed. To obtain it, further modeling work is needed, to test observed and putative interactions and to determine their effects at the population level. From an experimental perspective, models can be used to analyze available surveillance and experimental data, generate hypotheses regarding interaction mechanisms at play in transmission or infection, and test their likelihoods. Competing assumptions on the biological interaction processes can be assessed and the strength of interactions can also be estimated. From a public health viewpoint, these new models will help better estimate the burden of influenza virus interactions in terms of morbidity and mortality, the cost-effectiveness of interventions, and, critically, more accurately predict the real impact of control measures.

## Challenges

Integrating transmission and infection by multiple pathogens into mathematical models poses several challenges. From a methodological perspective, modeling several pathogens with interrelated natural histories makes classical compartmental approaches more difficult. Individual-based frameworks are better adapted for this task, e.g., this approach could be used to investigate the effect of the interval between influenza infection and bacterial acquisition, which reportedly affects the risk of bacterial invasion [30,122]. Individual-based models are often more computationally intensive and can introduce new difficulties in terms of parameter estimation, requiring the design of new methods. Recent developments in statistical inference methods, like particle Markov chain Monte Carlo (pMCMC) or maximum likelihood estimation via iterated filtering (MIF) [123,124], now enable modelers to jointly fit complex population-based models to multiple types of data, thereby allowing more data, and more diverse types of data to inform the model parameters.

Epidemiological data represent the second major challenge. To date, modeling studies have been limited by the poor knowledge of respiratory viruses and bacteria circulating in the community, especially because little is known about prevalence, incidence, at-risk populations and even epidemic profiles in different populations. On an individual level, new studies are required to assess the effect of coinfections, rather than ecological associations from incidence data. Important features include: i) coinfection-induced alteration of diseases’ natural histories, e.g. increased acquisition and severity risk, changes of infection durations and generation times; ii) specific at risk-periods for secondary infection or invasion of the coinfecting pathogen, or at-risk periods for severe outcomes; iii) at-risk populations, as characterized by individuals’ age, comorbidities or behavioral risk factors.

For population-level data, in most countries surveillance of influenza acquisition is based on networks of general practitioners who notify patients consulting for clinical symptoms of ILI [125]. Surveillance-data streams based on syndromic surveillance [126], inpatient data [127] and pathogen testing [128,129] should be combined, and linked at the patient level, to better identify non-influenza infections, or anomalous epidemics that could signal interaction. Improvement of data quality in patient records and detection of the biases inherent in different types of surveillance data are critical to achieve this goal. The latter could be reached by developing new microbiological tools, including new sampling kits able to rapidly detect multiple pathogens for use during medical consultations.

## Conclusion

We examined the epidemiological and biological evidence supporting influenza virus interference and interaction with other pathogens. We highlighted opportunities to improve knowledge and control of the virus, if we can move forward from the tunnel vision of singlepathogen models. It is time to develop a more holistic approach to pathogen dynamics in mathematical modeling, with novel methodological innovations, and further efforts in data collection and surveillance. The motivation to do so lies in the real opportunity to improve public health practices, and create better, more cost-effective interventions against influenza.

## Acknowledgments

The authors thank Elizabeth Miller and Matthieu Domenech de Celles for helpful comments on the manuscript. They thank the three anonymous reviewers for helpful and constructive comments. Box 1. Mathematical modeling definitions

### Box 1. Mathematical modeling definitions

It can be difficult to navigate studies using mathematical modelling for infectious diseases, because modelers use their own lexicon, and words depart from their colloquial meanings. Multiple words can also be used to describe the same thing (reflecting the multi-disciplinary roots of mathematical modeling).

*Mathematical vs statistical models:* a **mathematical model** (or **transmission** or **mechanistic model**) is a mechanistic description by mathematical equations of how the number of infected entities changes over time. Depending on the scale of the model, entities can be cells, individuals, or groups of individuals (e.g. a household, a city). **Statistical models** do not include a mechanistic link between quantities, but only rely on a functional relationship, often in the form of a probability distribution.

*Individual-based model vs compartmental models:* **Individual-based models** (or **agent-based models**) include a description of the properties of each of the individuals in the studied population. In contrast, **compartmental models** group individuals with similar characteristics together into the compartments, and look at relationship between these compartments. The most famous compartmental in epidemiology is the SIR model, based on three compartments, Susceptible-Infectious-Recovered, which is the basis of most existing models. Compartmental models are easier to fit to data and interpret. Individual-based models are more flexible when it is important to integrate a wide range of characteristics of the population, but are comparatively slow to implement and run, and require good data on each characteristic that is modeled.

*Model fitting:* Models are built around a structure (the mechanisms), which is modulated by parameters which govern the rates of change between compartments, disease states, behaviors, etc. Historically, parameters have been estimated using results from studies published in the literature. In recent years, with the increased availability of epidemiological data, modelers try whenever possible to **fit the model to data** (also called **parameter inference** or **calibration**). For this they use algorithms that explore “parameter space” which is the set of all possible values for parameters, and retain sets of parameters that explain the observed data best. Fitting can be computationally intensive if the model includes many parameters. More efficient fitting algorithms allow fitting of more complex models and thus study of potentially more interaction mechanisms.

### Box 2. Benefits of Transmission Models

- **Allow causal relationships to be drawn from the data by testing hypotheses regarding interaction mechanisms:**

→ For example, using models to analyze the cellular dynamics observed in vivo in mouse coinfection experiments it is possible to design models of hypothesized immunological pathways and determine which most closely fits observed patterns [90].
- **Accurately evaluate the influenza burden**

→ For example, for year-to-year influenza epidemics have a different estimated reporting fraction. A model could be used to determine if coinfection or concurrent epidemics of other viruses are the reason for an increased (or decreased) probability of reporting infection.
- **Predict or project incidence of co-infections, including during pandemics**

→ For example, fitting multi-pathogen models to respiratory virus surveillance data would allow quantitative assessment of the hypothesis that during the 2009 pandemic influenza affected the timing of rhinovirus, RSV and influenza by competition [63,74].
- **Optimize prevention and control of influenza infections and their complications**

→ For example, a model of influenza and pneumococcal pneumonia could determine optimal target groups for pneumococcal vaccination, based on both the bacterial carriage rates in each age group, and the expected influenza vaccination rates in those age groups.
- **Estimate the costs and benefits of intervention strategies**

→ For example, a model based analysis of *in vitro* experimental data could allow assessment of the impact of early antiviral or antibiotic treatment on probability of pneumococcal invasion [91,92]. Combined with population it would be possible to assess the impact on secondary bacterial infections.

### Box 3. A simple model of interaction

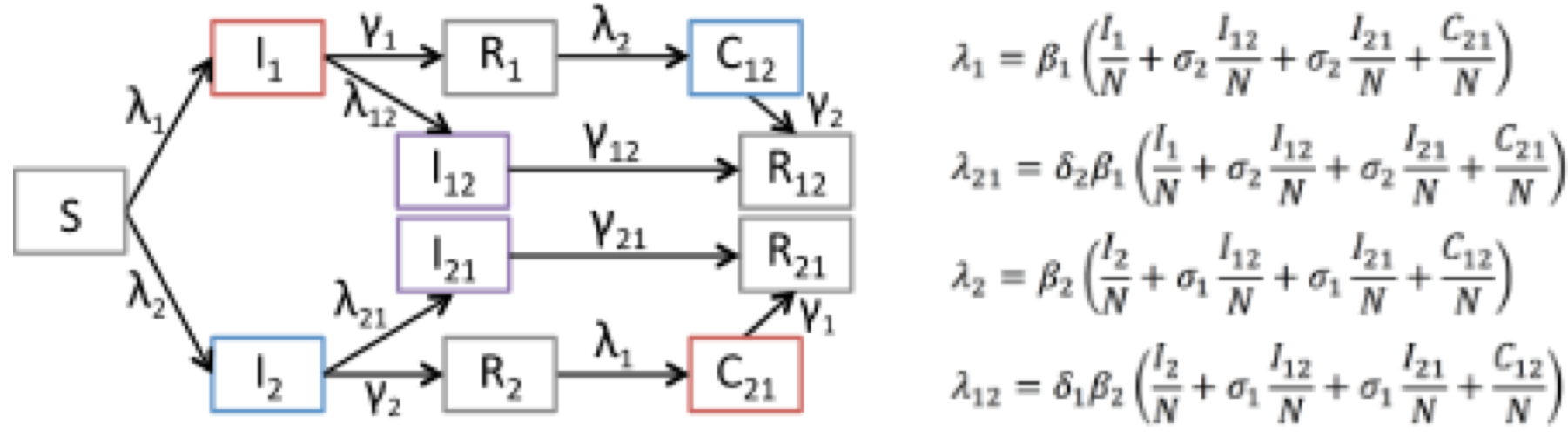

The simple model described here allows testing of two interaction mechanisms: increased (or decreased) infectiousness on coinfection, and decreased (or increased) probability of coinfection occurring. These are the two most commonly suggested mechanisms, the first of the “bacterial type” and the second of the “viral type” (Fig 3).

In the compartmental model figure above, all individuals start in the Susceptible (*S*) class, and move to the Infectious classes when they are infected by either pathogen 1 (*l*_*1*_ or 2 (*l*_*2*_).

Infected (and infectious) compartments are shown in color, where red is infectious with pathogen 1, blue marks infectious with pathogen 2, and infected and infectious with both pathogens in purple. Infection rates are given by the four forces of infection (*λ*_*1*_, *λ*_*2*_, *λ*_*12*_, *λ*_*21*_). After being infected by one pathogen, individuals can either be coinfected by the other pathogen and move to the coinfection compartments in purple (*l*_*12*_ or *l*_*21*_), or they can recover at rates *γ* and move to the Recovered compartments (*R*_*1*_ and *R*_*2*_). Coinfected individuals (*l*_*12*_ and *l*_*21*_) recover and remain in the doubly recovered compartments, *R*_*12*_ and *R*_*21*_. Individuals in *R*_*1*_ or *R*_*2*_ are subject to force of infection *λ*_*2*_ or *λ*_*1*_ respectively, i.e. of the pathogen they have not yet had. On infection with the other pathogen, they move to the consecutive infection compartment (*C*_*12*_ or *C*_*21*_). After recovery, those individuals move to the doubly recovered compartments (*R*_*12*_ and *R*_*21*_).

Parameters *β*_*1*_ and *β*_*2*_ are the baseline transmissibility of pathogen 1 and 2 respectively. There are four interaction parameters modulating the pathogen’s transmissibility: *σ*_*1*_ and *σ*_*2*_ are the change in infectiousness of coinfected classes, where a value less than 1 makes the coinfected class less infectious, and a value greater than 1 means coinfected individuals are more infectious. Parameters *δ*_*1*_ and *δ*_*2*_ alter the probability of acquisition of a second infection following a first infection, where a value less than 1 makes coinfection less likely, and a value above 1 makes it more likely.

Computer code generating these trajectories is given in Supplementary File S1.

## References

1. Webster RG, Monto, A. S., Braciale, T. J. & Lamb, R. A. (2013) Textbook of Influenza.

2. de Steenhuijsen Piters WAA, Sanders EAM, Bogaert D, Grice E, Segre J, et al. (2015) The role of the local microbial ecosystem in respiratory health and disease. Philosophical transactions of the Royal Society of London Series B, Biological sciences 370: 244–253.

3. Bosch AATM, Biesbroek G, Trzcinski K, Sanders EAM, Bogaert D (2013) Viral and bacterial interactions in the upper respiratory tract. PLoS pathogens 9: e1003057.

4. Yamin D, Balicer RD, Galvani AP (2014) Cost-effectiveness of influenza vaccination in prior pneumonia patients in Israel. Vaccine 32: 4198–4205.

5. Bedford T, Riley S, Barr IG, Broor S, Chadha M, et al. (2015) Global circulation patterns of seasonal influenza viruses vary with antigenic drift. Nature 523: 217–220.

6. Hayward AC, Fragaszy EB, Bermingham A, Wang L, Copas A, et al. (2014) Comparative community burden and severity of seasonal and pandemic influenza: results of the Flu Watch cohort study. Lancet Respir Med 2: 445–454.

7. de Francisco Shapovalova N, Donadel M, Jit M, Hutubessy R (2015) A systematic review of the social and economic burden of influenza in low- and middle-income countries. Vaccine 33: 6537–6544.

8. Brundage JF, Shanks GD (2008) Deaths from bacterial pneumonia during 1918-19 influenza pandemic. Emerging Infectious Diseases 14: 1193–1199.

9. Joseph C, Togawa Y, Shindo N (2013) Bacterial and viral infections associated with influenza. Influenza and Other Respiratory Viruses 7: 105–113.

10. Grassly NC, Fraser C (2008) Mathematical models of infectious disease transmission. Nat Rev Microbiol 6: 477–487.

11. Neher RA, Bedford T (2015) nextflu: real-time tracking of seasonal influenza virus evolution in humans. Bioinformatics 31: 3546–3548.

12. Viboud C, Bjornstad ON, Smith DL, Simonsen L, Miller MA, et al. (2006) Synchrony, waves, and spatial hierarchies in the spread of influenza. Science 312: 447–451.

13. Wu JT, Riley S, Fraser C, Leung GM (2006) Reducing the impact of the next influenza pandemic using household-based public health interventions. PLoS Med 3: e361.

14. Longini IM Jr., Halloran ME, Nizam A, Yang Y (2004) Containing pandemic influenza with antiviral agents. Am J Epidemiol 159: 623–633.

15. Gaglani MJ, Piedra PA, Herschler GB, Griffith ME, Kozinetz CA, et al. (2004) Direct and total effectiveness of the intranasal, live-attenuated, trivalent cold-adapted influenza virus vaccine against the 2000-2001 influenza A(H1N1) and B epidemic in healthy children. Arch Pediatr Adolesc Med 158: 65–73.

16. Matrajt L, Halloran ME, Longini IM, Jr., (2013) Optimal vaccine allocation for the early mitigation of pandemic influenza. PLoS Comput Biol 9: e1002964.

17. House T, Baguelin M, Van Hoek AJ, White PJ, Sadique Z, et al. (2011) Modelling the impact of local reactive school closures on critical care provision during an influenza pandemic. Proc Biol Sci 278: 2753–2760.

18. Ferguson NM, Cummings DA, Fraser C, Cajka JC, Cooley PC, et al. (2006) Strategies for mitigating an influenza pandemic. Nature 442: 448–452.

19. Baguelin M, Flasche S, Camacho A, Demiris N, Miller E, et al. (2013) Assessing optimal target populations for influenza vaccination programmes: an evidence synthesis and modelling study. PLoS Med 10: e1001527.

20. Opatowski L, Fraser C, Griffin J, de Silva E, Van Kerkhove MD, et al. (2011) Transmission characteristics of the 2009 H1N1 influenza pandemic: comparison of 8 Southern hemisphere countries. PLoS Pathog 7: e1002225.

21. Kucharski AJ, Kwok KO, Wei VW, Cowling BJ, Read JM, et al. (2014) The contribution of social behaviour to the transmission of influenza A in a human population. PLoS Pathog 10: e1004206.

22. Cauchemez S, Bhattarai A, Marchbanks TL, Fagan RP, Ostroff S, et al. (2011) Role of social networks in shaping disease transmission during a community outbreak of 2009 H1N1 pandemic influenza. Proc Natl Acad Sci U S A 108: 2825–2830.

23. Apolloni A, Poletto C, Colizza V (2013) Age-specific contacts and travel patterns in the spatial spread of 2009 H1N1 influenza pandemic. BMC Infect Dis 13: 176.

24. Shaman J, Pitzer VE, Viboud C, Grenfell BT, Lipsitch M (2010) Absolute humidity and the seasonal onset of influenza in the continental United States. PLoS Biol 8: e1000316.

25. Baguelin M, Hoek AJ, Jit M, Flasche S, White PJ, et al. (2010) Vaccination against pandemic influenza A/H1N1v in England: a real-time economic evaluation. Vaccine 28: 2370–2384.

26. Mina MJ, Klugman KP (2014) The role of influenza in the severity and transmission of respiratory bacterial disease. The Lancet Respiratory medicine 2: 750–763.

27. Short K, Habets M (2012) Interactions between Streptococcus pneumoniae and influenza virus: a mutually beneficial relationship? Future Microbiology: 609–624.

28. DaPalma T, Doonan BP, Trager NM, Kasman LM (2010) A systematic approach to virus-virus interactions. Virus research 149: 1–9.

29. Abramson JS, Mills EL, Giebink GS, Quie PG (1982) Depression of monocyte and polymorphonuclear leukocyte oxidative metabolism and bactericidal capacity by influenza A virus. Infection and immunity 35: 350–355.

30. Rynda-Apple A, Robinson KM, Alcorn JF (2015) Influenza and bacterial superinfection: Illuminating the immunologic mechanisms of disease. Infection and Immunity 83: 3764–3770.

31. Ghoneim HE, Thomas PG, McCullers Ja (2013) Depletion of alveolar macrophages during influenza infection facilitates bacterial superinfections. Journal of immunology (Baltimore, Md : 1950) 191: 1250–1259.

32. Sen GC (2001) Viruses and interferons. Annu Rev Microbiol 55: 255–281.

33. Loh E, Kugelberg E, Tracy A, Zhang Q, Gollan B, et al. (2013) Temperature triggers immune evasion by Neisseria meningitidis. Nature 502: 237–240.

34. Braun LE, Sutter DE, Eichelberger MC, Pletneva L, Kokai-Kun JF, et al. (2007) Co infection of the cotton rat (Sigmodon hispidus) with Staphylococcus aureus and influenza A virus results in synergistic disease. Microbial Pathogenesis 43: 208–216.

35. Smith AM, McCullers JA (2014) Secondary bacterial infections in influenza virus infection pathogenesis. Current topics in microbiology and immunology 385: 327–356.

36. Tashiro M, Ciborowski P, Reinacher M, Pulverer G, Klenk HD, et al. (1987) Synergistic role of staphylococcal proteases in the induction of influenza virus pathogenicity. Virology 157: 421–430.

37. Rohani P, Earn DJ, Finkenstädt B, Grenfell BT (1998) Population dynamic interference among childhood diseases. Proceedings Biological sciences / The Royal Society 265: 2033–2041.

38. Rohani P, Green CJ, Mantilla-Beniers NB, Grenfell BT (2003) Ecological interference between fatal diseases. Nature 422: 885–888.

39. Turner RD, Bothamley GH (2015) Cough and the transmission of tuberculosis. The Journal of infectious diseases 211: 1367–1372.

40. Cohen R, Varon E, Doit C, Schlemmer C, Romain O, et al. (2015) A 13-year survey of pneumococcal nasopharyngeal carriage in children with acute otitis media following PCV7 and PCV13 implementation. Vaccine 33: 5118–5126.

41. Yildirim I, Little BA, Finkelstein J, Lee G, Hanage WP, et al. (2017) Surveillance of pneumococcal colonization and invasive pneumococcal disease reveals shift in prevalent carriage serotypes in Massachusetts’ children to relatively low invasiveness. Vaccine 35: 4002–4009.

42. McCullers JA (2006) Insights into the interaction between influenza virus and pneumococcus. Clinical microbiology reviews 19: 571–582.

43. Zhou H, Haber M, Ray S, Farley MM, Panozzo CA, et al. (2012) Invasive pneumococcal pneumonia and respiratory virus co-infections. Emerg Infect Dis 18: 294–297.

44. Kim PE, Musher DM, Glezen WP, Rodriguez-Barradas MC, Nahm WK, et al. (1996) Association of invasive pneumococcal disease with season, atmospheric conditions, air pollution, and the isolation of respiratory viruses. Clin Infect Dis 22: 100–106.

45. Watson M, Gilmour R, Menzies R, Ferson M, McIntyre P, et al. (2006) The association of respiratory viruses, temperature, and other climatic parameters with the incidence of invasive pneumococcal disease in Sydney, Australia. Clin Infect Dis 42: 211–215.

46. Toschke AM, Arenz S, von Kries R, Puppe W, Weigl JA, et al. (2008) No temporal association between influenza outbreaks and invasive pneumococcal infections. Arch Dis Child 93: 218–220.

47. Sun K, Metzger DW (2014) Influenza Infection Suppresses NADPH Oxidase-Dependent Phagocytic Bacterial Clearance and Enhances Susceptibility to Secondary Methicillin-Resistant Staphylococcus aureus Infection. Journal of immunology (Baltimore, Md : 1950) 192: 3301–3307.

48. Lee LN, Dias P, Han D, Yoon S, Shea A, et al. (2010) A mouse model of lethal synergism between influenza virus and Haemophilus influenzae. The American journal of pathology 176: 800–811.

49. Hubert B, Watier L, Garnerin P, Richardson S (1992) Meningococcal disease and influenza-like syndrome: a new approach to an old question. The Journal of infectious diseases 166: 542–545.

50. Rameix-Welti M-A, Zarantonelli ML, Giorgini D, Ruckly C, Marasescu M, et al. (2009) Influenza A virus neuraminidase enhances meningococcal adhesion to epithelial cells through interaction with sialic acid-containing meningococcal capsules. Infection and immunity 77: 3588–3595.

51. Walaza S, Tempia S, Dawood H, Variava E, Moyes J, et al. (2015) Influenza virus infection is associated with increased risk of death amongst patients hospitalized with confirmed pulmonary tuberculosis in South Africa, 2010-2011. BMC Infectious Diseases 15: 26.

52. Flórido M, Grima Ma, Gillis CM, Xia Y, Turner SJ, et al. (2013) Influenza A virus infection impairs mycobacteria-specific T cell responses and mycobacterial clearance in the lung during pulmonary coinfection. J Immunol 191: 302–311.

53. Sonoguchi T, Naito H, Hara M, Takeuchi Y, Fukumi H (1985) Cross-Subtype Protection in Humans During Sequential, Overlapping, and/or Concurrent Epidemics Caused by H3N2 and H1N1 Influenza Viruses. Journal of Infectious Diseases 151: 81–88.

54. Laurie KL, Guarnaccia TA, Carolan LA, Yan AWC, Aban M, et al. (2015) Interval Between Infections and Viral Hierarchy Are Determinants of Viral Interference Following Influenza Virus Infection in a Ferret Model. Journal of Infectious Diseases 212: 1–10.

55. Bedford T, Rambaut A, Pascual M (2012) Canalization of the evolutionary trajectory of the human influenza virus. BMC biology 10: 38.

56. Boni MF, Gog JR, Andreasen V, Christiansen FB (2004) Influenza drift and epidemic size: the race between generating and escaping immunity. Theoretical population biology 65: 179–191.

57. Smith DJ, Lapedes AS, de Jong JC, Bestebroer TM, Rimmelzwaan GF, et al. (2004) Mapping the antigenic and genetic evolution of influenza virus. Science 305: 371–376.

58. Davenport FM, Hennessy AV, Francis T Jr., (1953) Epidemiologic and immunologic significance of age distribution of antibody to antigenic variants of influenza virus. J Exp Med 98: 641–656.

59. Gostic KM, Ambrose M, Worobey M, Lloyd-Smith JO (2016) Potent protection against H5N1 and H7N9 influenza via childhood hemagglutinin imprinting. Science 354: 722–726.

60. Kucharski AJ, Gog JR (2012) The role of social contacts and original antigenic sin in shaping the age pattern of immunity to seasonal influenza. PLoS Comput Biol 8: e1002741.

61. Skowronski DM, De Serres G, Crowcroft NS, Janjua NZ, Boulianne N, et al. (2010) Association between the 2008-09 Seasonal Influenza Vaccine and Pandemic H1N1 Illness during Spring-Summer 2009: Four Observational Studies from Canada. PLoS Medicine 7: e1000258.

62. Cowling BJ, Fang VJ, Nishiura H, Chan K-H, Ng S, et al. (2012) Increased risk of noninfluenza respiratory virus infections associated with receipt of inactivated influenza vaccine. Clinical infectious diseases : an official publication of the Infectious Diseases Society of America 54: 1778–1783.

63. Casalegno JS, Ottmann M, Duchamp MB, Escuret V, Billaud G, et al. (2010) Rhinoviruses delayed the circulation of the pandemic influenza A (H1N1) 2009 virus in France. Clinical microbiology and infection : the official publication of the European Society of Clinical Microbiology and Infectious Diseases 16: 326–329.

64. Yang L, Chan KH, Suen LKP, Chan KP, Wang X, et al. (2015) Impact of the 2009 H1N1 Pandemic on Age-Specific Epidemic Curves of Other Respiratory Viruses: A Comparison of Pre-Pandemic, Pandemic and Post-Pandemic Periods in a Subtropical City. PloS one 10: e0125447.

65. Navarro-Marí JM, Pérez-Ruiz M, Galán Montemayor JC, Marcos Maeso MÁ, Reina J, et al. (2012) Circulation of other respiratory viruses and viral co-infection during the 2009 pandemic influenza. Enfermedades infecciosas y microbiología clínica 30 Suppl 4: 25–31.

66. Yoshida L-M, Suzuki M, Nguyen HA, Le MN, Dinh Vu T, et al. (2013) Respiratory syncytial virus: co-infection and paediatric lower respiratory tract infections. The European respiratory journal 42: 461–469.

67. Richard N, Komurian-Pradel F, Javouhey E, Perret M, Rajoharison A, et al. (2008) The Impact of Dual Viral Infection in Infants Admitted to a Pediatric Intensive Care Unit Associated with Severe Bronchiolitis. The Pediatric Infectious Disease Journal 27: 213–217.

68. Marcos MA, Ramon S, Anton A, Martinez E, Vilella A, et al. (2011) Clinical relevance of mixed respiratory viral infections in adults with influenza A H1N1. The European respiratory journal 38: 739–742.

69. Schnepf N, Resche-Rigon M, Chaillon A, Scemla A, Gras G, et al. (2011) High burden of non-influenza viruses in influenza-like illness in the early weeks of H1N1v epidemic in France. PloS one 6: e23514.

70. Rhedin S, Hamrin J, Naucler P, Bennet R, Rotzén-Östlund M, et al. (2012) Respiratory Viruses in Hospitalized Children with Influenza-Like Illness during the H1n1 2009 Pandemic in Sweden. PLoS ONE 7: e51491.

71. Esper FP, Spahlinger T, Zhou L (2011) Rate and influence of respiratory virus co infection on pandemic (H1N1) influenza disease. The Journal of infection 63: 260–266.

72. Anestad G (1982) Interference between outbreaks of respiratory syncytial virus and influenza virus infection. Lancet (London, England) 1: 502.

73. Glezen WP, Paredes A, Taber LH (1980) Influenza in children. Relationship to other respiratory agents. JAMA 243: 1345–1349.

74. Casalegno JS, Ottmann M, Bouscambert-Duchamp M, Valette M, Morfin F, et al. (2010) Impact of the 2009 influenza A(H1N1) pandemic wave on the pattern of hibernal respiratory virus epidemics, France, 2009. Euro surveillance : bulletin Européen sur les maladies transmissibles = European communicable disease bulletin 15: 1.

75. Mak GC, Wong AH, Ho WYY, Lim W (2012) The impact of pandemic influenza A (H1N1) 2009 on the circulation of respiratory viruses 2009-2011. Influenza and other respiratory viruses 6: e6–10.

76. Yang Y, Wang Z, Ren L, Wang W, Vernet G, et al. (2012) Influenza A/H1N1 2009 pandemic and respiratory virus infections, Beijing, 2009-2010. PloS one 7: e45807.

77. Aberle JHMA, Stephan W. MD*; Pracher, Elisabeth MD‡; Hutter, Hans-Peter MD†; Kundi, Michael MD†; Popow-Kraupp, Therese MD* (2005) Single Versus Dual Respiratory Virus Infections in Hospitali…: The Pediatric Infectious Disease Journal. The pediatric infectious disease journal 24.

78. Goka E, Vallely P, Mutton K, Klapper P (2013) Influenza A viruses dual and multiple infections with other respiratory viruses and risk of hospitalisation and mortality. Influenza Other Respir Viruses 7: 1079–1087.

79. Walzl G, Tafuro S, Moss P, Openshaw PJ, Hussell T (2000) Influenza virus lung infection protects from respiratory syncytial virus-induced immunopathology. The Journal of experimental medicine 192: 1317–1326.

80. Pascalis H, Temmam S, Turpin M, Rollot O, Flahault A, et al. (2012) Intense Co Circulation of Non-Influenza Respiratory Viruses during the First Wave of Pandemic Influenza pH1N1/2009: A Cohort Study in Reunion Island. PLoS ONE 7: e44755.

81. Greer RM, McErlean P, Arden KE, Faux CE, Nitsche A, et al. (2009) Do rhinoviruses reduce the probability of viral co-detection during acute respiratory tract infections? Journal of Clinical Virology 45: 10–15.

82. Murphy BR, Richman DD, Chalhub EG, Uhlendorf CP, Baron S, et al. (1975) Failure of attenuated temperature-sensitive influenza A (H3N2) virus to induce heterologous interference in humans to parainfluenza type 1 virus. Infection and immunity 12: 62–68.

83. Nisii C, Meschi S, Selleri M, Bordi L, Castilletti C, et al. (2010) Frequency of detection of upper respiratory tract viruses in patients tested for pandemic H1N1/09 viral infection. Journal of clinical microbiology 48: 3383–3385.

84. Goka EA, Vallely PJ, Mutton KJ, Klapper PE (2014) Single and multiple respiratory virus infections and severity of respiratory disease: a systematic review. Paediatric respiratory reviews 15: 363–370.

85. Nicoli EJ, Trotter CL, Turner KME, Colijn C, Waight P, et al. (2013) Influenza and RSV make a modest contribution to invasive pneumococcal disease incidence in the UK. The Journal of infection 66: 512–520.

86. Weinberger DM, Harboe ZB, Krause TG, Miller M, Konradsen HB (2013) Serotype-specific effect of influenza on adult invasive pneumococcal pneumonia. Journal of Infectious Diseases: 1–22.

87. Gilca R, De Serres G, Skowronski D, Boivin G, Buckeridge DL (2009) The need for validation of statistical methods for estimating respiratory virus-attributable hospitalization. Am J Epidemiol 170: 925–936.

88. Boianelli A, Nguyen VK, Ebensen T, Schulze K, Wilk E, et al. (2015) Modeling Influenza Virus Infection: A Roadmap for Influenza Research. Viruses 7: 5274–5304.

89. Hodgson D, Baguelin M, van Leeuwen E, Panovska-Griffiths J, Ramsay M, et al. (2017) Effect of mass paediatric influenza vaccination on existing influenza vaccination programmes in England and Wales: a modelling and cost-effectiveness analysis. Lancet Public Health 2: e74–e81.

90. Smith AM, Adler FR, Ribeiro RM, Gutenkunst RN, McAuley JL, et al. (2013) Kinetics of coinfection with influenza A virus and Streptococcus pneumoniae. PLoS pathogens 9: e1003238.

91. Shrestha S, Foxman B, Dawid S, Aiello AE, Davis BM, et al. (2013) Time and dose-dependent risk of pneumococcal pneumonia following influenza: a model for within- host interaction between influenza and Streptococcus pneumoniae. Journal of the Royal Society, Interface / the Royal Society 10: 20130233.

92. Smith AM (2017) Quantifying the therapeutic requirements and potential for combination therapy to prevent bacterial coinfection during influenza. J Pharmacokinet Pharmacodyn 44: 81–93.

93. Boianelli A, Sharma-Chawla N, Bruder D, Hernandez-Vargas EA (2016) Oseltamivir PK/PD Modeling and Simulation to Evaluate Treatment Strategies against Influenza-Pneumococcus Coinfection. Front Cell Infect Microbiol 6: 60.

94. Smith AM, Smith AP (2016) A Critical, Nonlinear Threshold Dictates Bacterial Invasion and Initial Kinetics During Influenza. Sci Rep 6: 38703.

95. Opatowski L, Varon E, Dupont C, Temime L, van der Werf S, et al. (2013) Assessing pneumococcal meningitis association with viral respiratory infections and antibiotics: insights from statistical and mathematical models. Proceedings Biological sciences / The Royal Society 280: 20130519.

96. Shrestha S, Foxman B, Berus J, van Panhuis WG, Steiner C, et al. (2015) The role of influenza in the epidemiology of pneumonia. Scientific Reports 5: 15314.

97. Chien YW, Levin BR, Klugman KP (2012) The anticipated severity of a “1918-like” influenza pandemic in contemporary populations: The contribution of antibacterial interventions. PLoS ONE 7.

98. Crowe S, Utley M, Walker G, Grove P, Pagel C (2011) A model to evaluate mass vaccination against pneumococcus as a countermeasure against pandemic influenza. Vaccine 29: 5065–5077.

99. Handel A, Longini IM, Antia R (2009) Intervention strategies for an influenza pandemic taking into account secondary bacterial infections. Epidemics 1: 185–195.

100. Shrestha S, Foxman B, Weinberger DM, Steiner C, Viboud C, et al. (2013) Identifying the Interaction Between Influenza and Pneumococcal Pneumonia Using Incidence Data. Science Translational Medicine 5: 191ra184–191ra184.

101. Arduin H, Domenech de Celles M, Guillemot D, Watier L, Opatowski L (2017) An agent-based model simulation of influenza interactions at the host level: insight into the influenza-related burden of pneumococcal infections. BMC Infect Dis 17: 382.

102. Yan AWC, Cao P, Heffernan JM, McVernon J, Quinn KM, et al. (2017) Modelling cross reactivity and memory in the cellular adaptive immune response to influenza infection in the host. Journal of Theoretical Biology 413: 34–49.

103. Cao P, Yan AWC, Heffernan JM, Petrie S, Moss RG, et al. (2015) Innate Immunity and the Inter-exposure Interval Determine the Dynamics of Secondary Influenza Virus Infection and Explain Observed Viral Hierarchies. PLOS Computational Biology 11: e1004334.

104. Zakikhany K, Degail MA, Lamagni T, Waight P, Guy R, et al. (2011) Increase in invasive streptococcus pyogenes and streptococcus pneumoniae infections in England, December 2010 to January 2011. Eurosurveillance 16: 1–4.

105. Pinky L, Dobrovolny HM (2016) Coinfections of the Respiratory Tract: Viral Competition for Resources. PloS one 11: e0155589.

106. Kucharski AJ, Andreasen V, Gog JR (2016) Capturing the dynamics of pathogens with many strains. J Math Biol 72: 1–24.

107. Ferguson NM, Galvani AP, Bush RM (2003) Ecological and immunological determinants of influenza evolution. Nature 422: 428–433.

108. Ackerman E, Longini IM, Seaholm SK, Hedin ÅS (1990) Simulation of Mechanisms of Viral Interference in Influenza. International Journal of Epidemiology 19: 444–454.

109. Merler S, Poletti P, Ajelli M, Caprile B, Manfredi P (2008) Coinfection can trigger multiple pandemic waves. Journal of Theoretical Biology 254: 499–507.

110. Velasco-Hernández JX, Núñez-López M, Comas-García A, Cherpitel DEN, Ocampo MC, et al. (2015) Superinfection between Influenza and RSV Alternating Patterns in San Luis Potosí State, México. PLOS ONE 10: e0115674.

111. Araz OM, Galvani A, Meyers LA (2012) Geographic prioritization of distributing pandemic influenza vaccines. Health Care Management Science 15: 175–187.

112. Madhi Sa, Klugman KP (2004) A role for Streptococcus pneumoniae in virus-associated pneumonia. Nature medicine 10: 811–813.

113. Simonsen L, Taylor RJ, Young-Xu Y, Haber M, May L, et al. (2011) Impact of pneumococcal conjugate vaccination of infants on pneumonia and influenza hospitalization and mortality in all age groups in the United States. MBio 2: e00309–00310.

114. Tsai Y-H, Hsieh M-J, Chang C-J, Wen Y-W, Hu H-C, et al. (2015) The 23-valent pneumococcal polysaccharide vaccine is effective in elderly adults over 75 years old—Taiwan's PPV vaccination program. Vaccine 33: 2897–2902.

115. Fleming-Dutra KE, Hersh AL, Shapiro DJ, Bartoces M, Enns EA, et al. (2016) Prevalence of Inappropriate Antibiotic Prescriptions Among US Ambulatory Care Visits, 2010-2011. JAMA 315: 1864.

116. Polgreen PM, Yang M, Laxminarayan R, Cavanaugh JE (2011) Respiratory fluoroquinolone use and influenza. Infection control and hospital epidemiology 32: 706–709.

117. Mina MJ, Klugman KP, McCullers JA (2013) Live Attenuated Influenza Vaccine, But Not Pneumococcal Conjugate Vaccine, Protects Against Increased Density and Duration of Pneumococcal Carriage After Influenza Infection in Pneumococcal Colonized Mice. Journal of Infectious Diseases 208: 1281–1285.

118. Muthuri SG, Venkatesan S, Myles PR, Leonardi-Bee J, Al Khuwaitir TSA, et al. (2014) Effectiveness of neuraminidase inhibitors in reducing mortality in patients admitted to hospital with influenza A H1N1pdm09 virus infection: a meta-analysis of individual participant data. The Lancet Respiratory medicine 2: 395–404.

119. Fry AM (2014) Effectiveness of neuraminidase inhibitors for severe influenza. The Lancet Respiratory Medicine. pp. 346–348.

120. McCullers Ja (2014) The public health policy implications of understanding metabiosis. Cell host & microbe 16: 3–4.

121. McCullers JA (2011) Preventing and treating secondary bacterial infections with antiviral agents. Antivir Ther 16: 123–135.

122. McCullers Ja, Rehg JE (2002) Lethal synergism between influenza virus and Streptococcus pneumoniae: characterization of a mouse model and the role of platelet-activating factor receptor. The Journal of infectious diseases 186: 341–350.

123. Andrieu C, Doucet A, Holenstein R (2010) Particle Markov chain Monte Carlo methods. Journal of the Royal Statistical Society Series B-Statistical Methodology 72: 269–342.

124. King AA, Nguyen D, Ionides EL (2016) Statistical Inference for Partially Observed Markov Processes via the R Package pomp. Journal of Statistical Software 69.

125. Monto AS (2002) Epidemiology of viral respiratory infections. The American Journal of Medicine 112: 4–12.

126. Stewart M, Loschen W, Kass-Hout T Enabling ESSENCE to Process and Export Meaningful Use Syndromic Surveillance Data.

127. Warren-Gash C, Bhaskaran K, Hayward A, Leung GM, Lo S-V, et al. (2011) Circulating influenza virus, climatic factors, and acute myocardial infarction: a time series study in England and Wales and Hong Kong. The Journal of infectious diseases 203: 1710–1718.

128. Mustaquim D (2014) The Evolution of the WHO/NREVSS Influenza Surveillance System: The Challenges and Opportunities that Accompany Electronic Laboratory Data. Online Journal of Public Health Informatics 6.

129. Zhao H, Green H, Lackenby A, Donati M, Ellis J, et al. (2014) A new laboratory-based surveillance system (Respiratory DataMart System) for influenza and other respiratory viruses in England: results and experience from 2009 to 2012. Eurosurveillance 19: 20680.

130. Wolf AI, Strauman MC, Mozdzanowska K, Whittle JRR, Williams KL, et al. (2014) Coinfection with Streptococcus pneumoniae Modulates the B Cell Response to Influenza Virus. Journal of Virology 88: 11995–12005.

131. Siegel Steven J, Roche Aoife M, Weiser Jeffrey N (2014) Influenza Promotes Pneumococcal Growth during Coinfection by Providing Host Sialylated Substrates as a Nutrient Source. Cell Host & Microbe 16: 55–67.

132. McCullers Ja, McAuley JL, Browall S, Iverson AR, Boyd KL, et al. (2010) Influenza enhances susceptibility to natural acquisition of and disease due to Streptococcus pneumoniae in ferrets. The Journal of infectious diseases 202: 1287–1295.

133. Peltola VT, Boyd KL, McAuley JL, Rehg JE, McCullers JA (2006) Bacterial sinusitis and otitis media following influenza virus infection in ferrets. Infection and immunity 74: 2562–2567.

134. Walters K-A, D'Agnillo F, Sheng Z-M, Kindrachuk J, Schwartzman LM, et al. (2016) 1918 pandemic influenza virus and Streptococcus pneumoniae co-infection results in activation of coagulation and widespread pulmonary thrombosis in mice and humans. The Journal of Pathology 238: 85–97.

135. Nakamura S, Davis KM, Weiser JN, Bogaert D, Groot RD, et al. (2011) Synergistic stimulation of type I interferons during influenza virus coinfection promotes Streptococcus pneumoniae colonization in mice. Journal of Clinical Investigation 121: 3657–3665.

136. Walter ND, Taylor TH, Shay DK, Thompson WW, Brammer L, et al. (2010) Influenza circulation and the burden of invasive pneumococcal pneumonia during a nonpandemic period in the United States. Clinical infectious diseases : an official publication of the Infectious Diseases Society of America 50: 175–183.

137. Nelson GE, Gershman Ka, Swerdlow DL, Beall BW, Moore MR (2012) Invasive pneumococcal disease and pandemic (H1N1) 2009, Denver, Colorado, USA. Emerging infectious diseases 18: 208–216.

138. Jansen AGSC, Sanders EAM, VAN DER Ende A, VAN Loon AM, Hoes AW, et al. (2008) Invasive pneumococcal and meningococcal disease: association with influenza virus and respiratory syncytial virus activity? Epidemiology and infection 136: 1448–1454.

139. Kuster SP, Tuite AR, Kwong JC, McGeer A, Fisman DN (2011) Evaluation of coseasonality of influenza and invasive pneumococcal disease: results from prospective surveillance. PLoS medicine 8: e1001042.

140. Ampofo K, Bender J, Sheng X, Korgenski K, Daly J, et al. (2008) Seasonal invasive pneumococcal disease in children: role of preceding respiratory viral infection. Pediatrics 122: 229–237.

141. Grabowska K, Hogberg L, Penttinen P, Svensson A, Ekdahl K (2006) Occurrence of invasive pneumococcal disease and number of excess cases due to influenza. BMC Infect Dis 6: 58.

142. Murdoch DR, Jennings LC (2009) Association of respiratory virus activity and environmental factors with the incidence of invasive pneumococcal disease. J Infect 58: 37–46.

143. Edwards LJ, Markey PG, Cook HM, Trauer JM, Krause VL (2011) The relationship between influenza and invasive pneumococcal disease in the Northern Territory, 2005-2009. Med J Aust 194: 207.

144. Weinberger DM, Harboe ZB, Viboud C, Krause TG, Miller M, et al. (2014) Pneumococcal disease seasonality: incidence, severity and the role of influenza activity. Eur Respir J 43: 833–841.

145. Grijalva CG, Griffin MR, Edwards KM, Williams JV, Gil AI, et al. (2014) The role of influenza and parainfluenza infections in nasopharyngeal pneumococcal acquisition among young children. Clin Infect Dis 58: 1369–1376.

146. Zhou H, Haber M, Ray S, Farley MM, Panozzo CA, et al. (2012) Invasive pneumococcal pneumonia and respiratory virus co-infections. Emerging infectious diseases 18: 294–297.

147. Damasio GAC, Pereira LA, Moreira SDR, Duarte dos Santos CN, Dalla-Costa LM, et al. (2015) Does virus-bacteria coinfection increase the clinical severity of acute respiratory infection? Journal of Medical Virology 87: 1456–1461.

148. Hendriks W, Boshuizen H, Dekkers A, Knol M, Donker GA, et al. (2017) Temporal cross-correlation between influenza-like illnesses and invasive pneumococcal disease in The Netherlands. Influenza Other Respir Viruses 11: 130–137.

149. Niemann S, Ehrhardt C, Medina E, Warnking K, Tuchscherr L, et al. (2012) Combined action of influenza virus and Staphylococcus aureus panton-valentine leukocidin provokes severe lung epithelium damage. The Journal of infectious diseases 206: 1138–1148.

150. Davison VE, Sanford BA (1982) Factors influencing adherence of Staphylococcus aureus to influenza A virus-infected cell cultures. Infection and Immunity 37: 946–955.

151. Zhang WJ, Sarawar S, Nguyen P, Daly K, Rehg JE, et al. (1996) Lethal synergism between influenza infection and staphylococcal enterotoxin B in mice. J Immunol 157: 5049–5060.

152. Chertow DS, Kindrachuk J, Sheng Z-M, Pujanauski LM, Cooper K, et al. (2016) Influenza A and Methicillin-resistant Staphylococcus aureus Co-infection in Rhesus Macaques -- A Model of Severe Pneumonia. Antiviral Research.

153. Iverson AR, Boyd KL, McAuley JL, Plano LR, Hart ME, et al. (2011) Influenza virus primes mice for pneumonia from Staphylococcus aureus. The Journal of infectious diseases 203: 880–888.

154. Robinson KM, Choi SM, McHugh KJ, Mandalapu S, Enelow RI, et al. (2013) Influenza A Exacerbates Staphylococcus aureus Pneumonia by Attenuating IL-1 Production in Mice. The Journal of Immunology 191: 5153–5159.

155. Sherertz RJ, Reagan DR, Hampton KD, Robertson KL, Streed SA, et al. (1996) A cloud adult: the Staphylococcus aureus-virus interaction revisited. Ann Intern Med 124: 539–547.

156. Hageman JC, Uyeki TM, Francis JS, Jernigan DB, Wheeler JG, et al. (2006) Severe community-acquired pneumonia due to Staphylococcus aureus, 2003-04 influenza season. Emerging infectious diseases 12: 894–899.

157. Finelli L, Fiore A, Dhara R, Brammer L, Shay DK, et al. (2008) Influenza-associated pediatric mortality in the United States: increase of Staphylococcus aureus coinfection. Pediatrics 122: 805–811.

158. Reed C, Kallen AJ, Patton M, Arnold KE, Farley MM, et al. (2009) Infection With Community-Onset Staphylococcus aureus and Influenza Virus in Hospitalized Children. The Pediatric Infectious Disease Journal 28: 572–576.

159. Kobayashi SD, Olsen RJ, LaCasse RA, Safronetz D, Ashraf M, et al. (2013) Seasonal H3N2 influenza A virus fails to enhance Staphylococcus aureus co-infection in a non-human primate respiratory tract infection model. Virulence 4: 707–715.

160. Michaels RH, Myerowitz RL, Klaw R (1977) Potentiation of experimental meningitis due to Haemophilus influenzae by influenza A virus. The Journal of infectious diseases 135: 641–645.

161. Bakaletz LO, Hoepf TM, Demaria TF, Lim DJ (1988) The Effect of Antecedent Influenza A Virus Infection on the Adherence of Hemophilus Influenzae to Chinchilla Tracheal Epithelium. Am ~ Otalaryngol 9: 127–134.

162. Francis T, De Torregrosa MV (1945) Combined infection of mice with H. Influenzae and influenza virus by the intranasal route. Journal of Infectious Diseases 76: 70–77.

163. Morens DM, Taubenberger JK, Fauci AS (2008) Predominant role of bacterial pneumonia as a cause of death in pandemic influenza: implications for pandemic influenza preparedness. The Journal of infectious diseases 198: 962–970.

164. Read RC, Goodwin L, Parsons MA, Silcocks P, Kaczmarski EB, et al. (1999) Coinfection with influenza B virus does not affect association of Neisseria meningitidis with human nasopharyngeal mucosa in organ culture. Infection and immunity 67: 3082–3086.

165. Cartwright KA, Jones DM, Smith AJ, Stuart JM, Kaczmarski EB, et al. (1991) Influenza A and meningococcal disease. Lancet (London, England) 338: 554–557.

166. Jacobs JH, Viboud C, Tchetgen ET, Schwartz J, Steiner C, et al. (2014) The Association of Meningococcal Disease with Influenza in the United States, 19892009. PloS one 9: e107486.

167. Brundage JF (2006) Interactions between influenza and bacterial respiratory pathogens: implications for pandemic preparedness. The Lancet Infectious diseases 6: 303–312.

168. Makras P, Alexiou-Daniel S, Antoniadis A, Hatzigeorgiou D (2001) Outbreak of meningococcal disease after an influenza B epidemic at a Hellenic Air Force recruit training center. Clinical infectious diseases : an official publication of the Infectious Diseases Society of America 33: e48–50.

169. Florido M, Pillay R, Gillis CM, Xia Y, Turner SJ, et al. (2015) Epitope-specific CD4+, but not CD8+, T-cell responses induced by recombinant influenza A viruses protect against Mycobacterium tuberculosis infection. European Journal of Immunology 45: 780–793.

170. Volkert M, Pierce C, Horsfall FL, Dubos RJ (1947) The enhancing effect of concurrent infection with pneumotropic viruses on pulmonary tuberculosis in mice. The Journal of experimental medicine 86: 203–214.

171. Redford PS, Mayer-Barber KD, McNab FW, Stavropoulos E, Wack A, et al. (2014) Influenza A virus impairs control of Mycobacterium tuberculosis coinfection through a type I interferon receptor-dependent pathway. The Journal of infectious diseases 209: 270–274.

172. Walaza S, Cohen C, Nanoo A, Cohen AL, McAnerney J, et al. (2015) Excess mortality associated with influenza among tuberculosis deaths in South Africa, 1999-2009. PLoS ONE 10: 1999–2009.

173. Oei W, Nishiura H, Oei W, Nishiura H (2012) The Relationship between Tuberculosis and Influenza Death during the Influenza (H1N1) Pandemic from 1918-19. Computational and Mathematical Methods in Medicine 2012: 1–9.

174. Noymer A (2011) The 1918 influenza pandemic hastened the decline of tuberculosis in the United States: An age, period, cohort analysis. Vaccine 29: B38–B41.

175. Noymer A (2009) Testing the influenza-tuberculosis selective mortality hypothesis with Union Army data. Social Science & Medicine 68: 1599–1608.

176. Zörcher K, Zwahlen M, Ballif M, Rieder HL, Egger M, et al. (2016) Influenza Pandemics and Tuberculosis Mortality in 1889 and 1918: Analysis of Historical Data from Switzerland. PLOS ONE 11: e0162575.

177. Roth S, Whitehead S, Thamthitiwat S, Chittaganpitch M, Maloney SA, et al. (2013) Concurrent influenza virus infection and tuberculosis in patients hospitalized with respiratory illness in Thailand. Influenza and other Respiratory Viruses 7: 244–248.

178. Klonoski JM, Hurtig HR, Juber BA, Schuneman MJ, Bickett TE, et al. (2014) Vaccination against the M protein of Streptococcus pyogenes prevents death after influenza virus:S. pyogenes super-infection. Vaccine 32: 5241–5249.

179. Okamoto S, Kawabata S, Nakagawa I, Okuno Y, Goto T, et al. (2003) Influenza A Virus-Infected Hosts Boost an Invasive Type of Streptococcus pyogenes Infection in Mice. Journal of Virology 77: 4104–4112.

180. Okamoto S, Kawabata S, Terao Y, Fujitaka H, Okuno Y, et al. (2004) The Streptococcus pyogenes Capsule Is Required for Adhesion of Bacteria to Virus-Infected Alveolar Epithelial Cells and Lethal Bacterial-Viral Superinfection. Infection and Immunity 72: 6068–6075.

181. Hafez MM, Abdel-Wahab KSE, El-Fouhil DFI (2010) Augmented adherence and internalization of group A Streptococcus pyogenes to influenza A virus infected MDCK cells. Journal of Basic Microbiology 50: S46–S57.

182. Scaber J, Saeed S, Ihekweazu C, Efstratiou A, Mccarthy N, et al. (2011) Group A streptococcal infections during the seasonal influenza outbreak 2010/11 in South East England. Euro Surveill 16.

183. Tasher D, Stein M, Simões EAF, Shohat T, Bromberg M, et al. (2011) Invasive bacterial infections in relation to influenza outbreaks, 2006-2010. Clinical infectious diseases : an official publication of the Infectious Diseases Society of America 53: 1199–1207.

184. Tamayo E, Montes M, Vicente D, Perez-Trallero E, Welch C, et al. (2016) Streptococcus pyogenes Pneumonia in Adults: Clinical Presentation and Molecular Characterization of Isolates 2006-2015. PLOS ONE 11: e0152640.

185. Anestad G, Vainio K, Hungnes O (2007) Interference between outbreaks of epidemic viruses. Scandinavian journal of infectious diseases 39: 653–654.

186. Ånestad G (2009) Surveillance of respiratory viral infections by rapid immunofluorescence diagnosis, with emphasis on virus interference. Epidemiology and Infection 99: 523.

187. Anestad G (1987) Surveillance of respiratory viral infections by rapid immunofluorescence diagnosis, with emphasis on virus interference. Epidemiol Infect 99: 523–531.

188. Nishimura N, Nishio H, Lee MJ, Uemura K (2005) The clinical features of respiratory syncytial virus: lower respiratory tract infection after upper respiratory tract infection due to influenza virus. Pediatr Int 47: 412–416.

189. van Asten L, Bijkerk P, Fanoy E, van Ginkel A, Suijkerbuijk A, et al. (2016) Early occurrence of influenza A epidemics coincided with changes in occurrence of other respiratory virus infections. Influenza and other respiratory viruses 10: 14–26.

190. Meningher T, Hindiyeh M, Regev L, Sherbany H, Mendelson E, et al. (2014) Relationships between A(H1N1)pdm09 influenza infection and infections with other respiratory viruses. Influenza and other respiratory viruses 8: 422–430.

191. Martin ET, Fairchok MP, Stednick ZJ, Kuypers J, Englund JA (2013) Epidemiology of multiple respiratory viruses in childcare attendees. The Journal of infectious diseases 207: 982–989.

192. Shinjoh M, Omoe K, Saito N, Matsuo N, Nerome K (2000) In vitro growth profiles of respiratory syncytial virus in the presence of influenza virus. Acta Virol 44: 91–97.

193. Linde A, Rotzen-Ostlund M, Zweygberg-Wirgart B, Rubinova S, Brytting M (2009) Does viral interference affect spread of influenza? Euro Surveill 14.

194. Anestad G, Nordbo SA (2011) Virus interference. Did rhinoviruses activity hamper the progress of the 2009 influenza A (H1N1) pandemic in Norway? Med Hypotheses 77: 1132–1134.

195. Tanner H, Boxall E, Osman H (2012) Respiratory viral infections during the 2009-2010 winter season in Central England, UK: incidence and patterns of multiple virus coinfections. European journal of clinical microbiology & infectious diseases : official publication of the European Society of Clinical Microbiology 31: 3001–3006.

196. Mackay IM, Lambert SB, Faux CE, Arden KE, Nissen MD, et al. (2013) Community-wide, contemporaneous circulation of a broad spectrum of human rhinoviruses in healthy Australian preschool-aged children during a 12-month period. The Journal of infectious diseases 207: 1433–1441.

197. Easton AJ, Scott PD, Edworthy NL, Meng B, Marriott AC, et al. (2011) A novel broad-spectrum treatment for respiratory virus infections: influenza-based defective interfering virus provides protection against pneumovirus infection in vivo. Vaccine 29: 2777–2784.

198. Goto H, Ihira H, Morishita K, Tsuchiya M, Ohta K, et al. (2016) Enhanced growth of influenza A virus by coinfection with human parainfluenza virus type 2. Med Microbiol Immunol 205: 209–218.

